# Time calibrated morpho-molecular classification of Nassellaria (Radiolaria)

**DOI:** 10.1101/435123

**Authors:** Miguel M. Sandin, Loic Pillet, Tristan Biard, Camille Poirier, Estelle Bigeard, Sarah Romac, Noritoshi Suzuki, Fabrice Not

**Affiliations:** Sorbonne Université, CNRS - UMR7144 - Ecology of Marine Plankton Group - Station Biologique de Roscoff, 29680 Roscoff, France; Department of Earth Science, Graduate School of Science, Tohoku University, Sendai 980–8578, Japan

## Abstract

Nassellaria are marine radiolarian protists belonging to the Rhizaria lineage. Their skeleton, made of opaline silica, exhibit an excellent fossil record, extremely valuable in micro-paleontological studies for paleo-environmental reconstruction. Yet, to date very little is known about the extant diversity and ecology of Nassellaria in contemporary oceans, and most of it is inferred from their fossil record. Here we present an integrative classification of Nassellaria based on taxonomical marker genes (18S and 28S ribosomal DNA) and morphological characteristics obtained by optical and scanning electron microscopy imaging. Our phylogenetic analyses distinguished 11 main morpho-molecular clades relying essentially on the overall morphology of the skeleton and not on internal structures as previously considered. Using fossil calibrated molecular clock we estimated the origin of Nassellaria among radiolarians primitive forms in the Devonian (ca. 420 Ma), that gave rise to living nassellarian groups in the Triassic (ca. 250 Ma), during the biggest diversification event over their evolutionary history. This morpho-molecular framework provides both a new morphological classification easier to identify under light microscopy and the basis for future molecular ecology surveys. Altogether, it brings a new standpoint to improve our scarce understanding of the ecology and worldwide distribution of extant nassellarians.

## Introduction

Along with Foraminifera, Radiolaria constitute the Phylum Retaria, within the supergroup Rhizaria, one of the 8 major branches of eukaryotic life (Burki and Keeling, 2014). Radiolarians are marine heterotrophic protists, currently classified in 5 taxonomic orders based on morphological features and chemical composition of their biomineralized skeleton. Acantharia possess a skeleton made out of strontium sulphate (SrSO_4_), while opaline silica (SiO_2_ nH_2_O) is found in skeletons of Taxopodia and the polycystines Collodaria, Nassellaria and Spumellaria (Suzuki and Not, 2015). The robust silica skeleton of polycystines preserves well in sediments and hard sedimentary rocks, providing an extensive fossil record throughout the Phanerozoic (De Wever et al., 2001). Essentially studied by micro-palaeontologists, classification and evolutionary history of Radiolaria are largely based on morphological criteria (Suzuki and Oba, 2015) and very little is known about the ecology and diversity of contemporary species.

Among polycystines, Nassellaria actively feed on a large variety of preys, from bacteria to mollusc larvae (Anderson, 1993; Sugiyama et al., 2008), contributing significantly to trophic webs dynamic of oceanic ecosystems. Some nassellarian species host photosynthetic algal symbionts, up to 50 symbionts per host cells, mainly identified as dinoflagellates (Suzuki and Not, 2015; Zhang et al., 2018, Decelle et al., 2015; Probert et al., 2014). This mixotrophic behaviour may influence their distribution patterns, being in surface tropical waters the greatest diversity and abundance values, and decreasing towards the poles and at depth (Boltovskoy and Correa, 2016; Boltovskoy, 2017). Only a few taxa are restricted to deep waters (1000-3000m) in which no photosymbionts have ever been described (Suzuki and Not, 2015).

Unlike other radiolarians, nassellarian skeleton is heteropolar, aligned along an axis and not a centre. This skeleton is divided in three main different segments: the cephalis always present (or 1st segment), the thorax (2nd segment) and sometimes an abdomen (3rd segment) and post-abdominal segments (Campbell, 1954). The cephalis contains the initial spicular system, widely used for the taxonomic classification at family or higher levels (De Wever et al., 2001; Petrushevskaya, 1971b). Its basic architecture is the component of *A*-rod (apical rod), *D*- (dorsal rod), *V*- (ventral rod), *MB* (median bar), occasionally Ax (axobate node), *l*-rod (lateral rod from *MB* at the *A*-rod side) and *L*-rod (lateral rod from *MB* at the *V*-rod side). All these initial spicules except for *V*- and *l-*rods are always present in Nassellaria (e.g. Fig 14 in De Wever et al., 2001). The architecture of the initial spicular system has been used not only in nassellarian classification but also for Collodaria, Spumellaria and other Polycystines such as Entactinaria, a group considered to be an early lineage in the Paleozoic. Morphology based taxonomic classifications have divided extant Nassellaria in nearly 25 families, 140 genera and 430 recognized species (Suzuki and Not, 2015). At higher level, they are currently divided in 7 super-families: Acanthodesmoidea (Hertwig, 1879; sensu Dumitrica in De Wever et al., 2001), Acropyramioidea (Haeckel, 1882; sensu emend. Petrushevskaya, 1981), Artostrobioidea (Riedel, 1967; sensu O’Dogherty, 1994), Cannobotryoidea (Haeckel, 1882; sensu Petrushevskaya, 1971a), Eucyrtidioidea (Ehrenberg, 1846; sensu Dumitrica in De Wever et al., 2001), Plagiacanthoidea (Hertwig, 1879; sensu Petrushevskaya, 1971a), Pterocorythoidea (Haeckel, 1882; sensu Matsuzaki et al., 2015) and some undetermined families (e.g. Theopiliidae, Bekomidae, Carpocaniidae) (Matsuzaki et al., 2015). The super-family Acanthodesmoidea includes almost all monocyrtid nassellarians (skeleton consists only of the cephalis segment), two spines from their internal spicular system (apical and ventral rods) have merged together forming the so-called D-ring (Fig. 2.E), characteristic of this group (De Wever et al., 2001). The Acropyramioidea is composed by dicyrtid nassellarians (cephalis and thorax) with a reduced cephalis and a pyramidal shape skeleton (Fig. 2.B). Multicyrtid nassellarians (cephalis, thorax, abdomen and post-abdominal segments) with a simple cephalis and a small apical rod belong to the Eucyrtidioidea (Fig. 2.A) (De Wever et al., 2001, Matsuzaki et al., 2015) whereas those with a flattered and more complex head (e.g. cephalis) and a well-developed ventral tube to the Artostrobioidea (Fig. 2.D). The Plagiacanthoidea and Cannobotryoidea are composed by dicyrtid nassellarians with a large cephalis relative to the whole skeleton. The former superfamily has a big cephalis relative to their test size but not subdivided (Hertwig, 1879; Petrushevskaya, 1971a) (Fig. 2.G1, G2, G4-G6), whereas in the second the cephalis is subdivided in lobes (Haeckel, 1882; Petrushevskaya, 1971a) (Fig. 2.G3). The last Superfamily, the Pterocorythoidea, associates di- or tri-cyrtid nassellarians with an elongated or spherical head and a stout apical horn (Haeckel, 1882; Matsuzaki et al., 2015) (Fig. 2.I-J).

**Figure 1.**
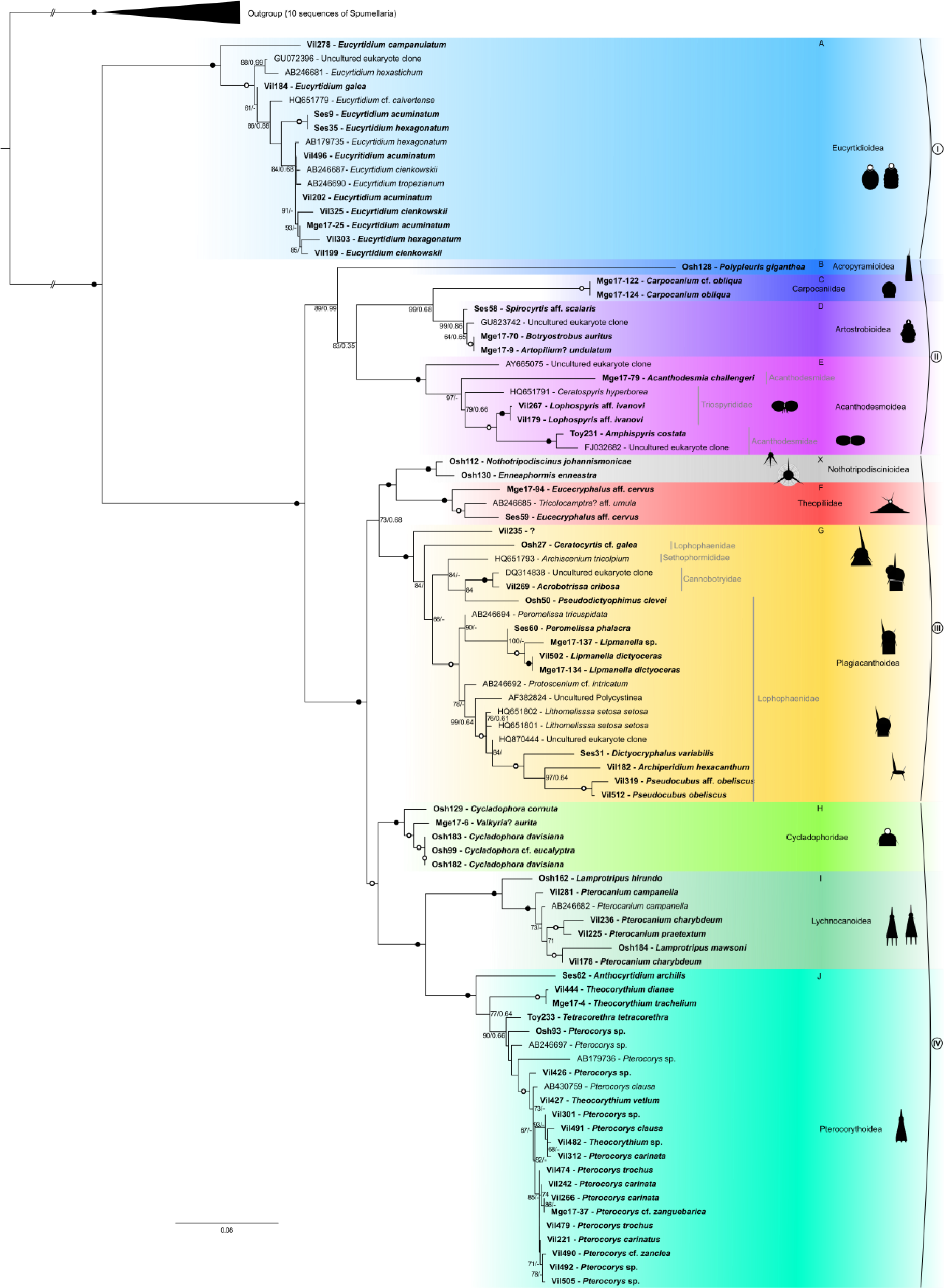
Molecular phylogeny of Nassellaria inferred from the concatenated complete 18S and partial 28S (D1-D2 regions) rDNA genes (90 taxa and 2590 aligned positions). The tree was obtained by using a phylogenetic Maximum likelihood method implemented using the GTR + γ + I model of sequence evolution. PhyML bootstrap values (100 000 replicates, BS) and posterior probabilities (PP) are shown at the nodes (BS/PP). Black circles indicate BS of 100% and PP > 0.99. Hollow circles indicates BS > 90% and PP > 0.90. Sequences obtained in this study are shown in bold. Eleven main clades are defined based on statistical support and morphological criteria (A, B, C, D, E, X, F, G, H, I, J). Figures besides clade names represent main features in the overall morphology of specimens included in the phylogeny. Ten Spumellaria sequences were assembled as out-group. Branches with a double barred symbol are fourfold reduced for clarity.

**Figure 2.**
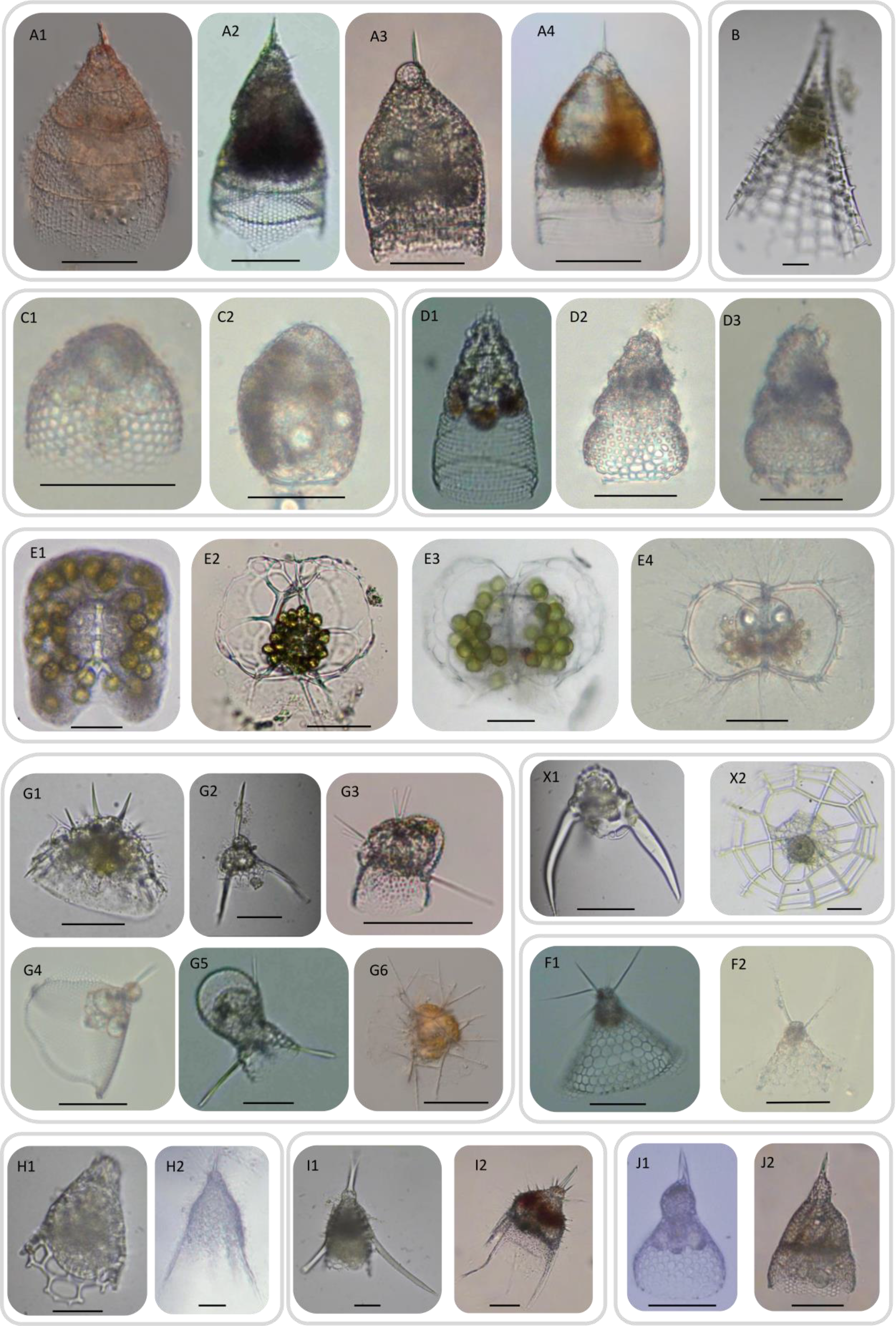
Light Microscopy (LM) images of live Nassellarian specimens used in this study for phylogenetic analysis. Letters correspond to its phylogenetic clade in Fig. 1. Scale bars (when available) = 50μm. (**A1**) Vil496: *Eucyritidium acuminatum*. (**A2**) Ses35: *Eucyrtidium hexagonatum*. (**A3**) Vil278: *Eucyrtidium campanulatum*. (**A4**) Vil184: *Eucyrtidium galea*. (**B**) Osh128: *Polypleuris giganthea*. (**C1**) Mge17-122: *Carpocanium* cf. *obliqua*. (**C2**) Mge17-124: *Carpocanium obliqua*. (**D1**) Ses58: *Spirocyrtis* aff. *Scalaris*. (**D2**) Mge17-9: *Artopilium*? *undulatum*. (**D3**) Mge17-70: *Botryostrobus auritus*. (**E1**) Toy231: *Amphispyris costata*. (**E2**) Vil267: *Lophospyris* aff*. ivanovi*. (**E3**) Vil179: *Lophospyris* aff*. ivanovi*. (**E4**) Mge17-79: *Acanthodesmia challengeri*. (**X1**) Osh112: *Nothotripodiscinus johannismonicae*. (**X2**) Osh130: *Enneaphormis enneastra.* (**F1**) Ses59: *Eucecryphalus* aff*. cervus*. (**F2**) Mge17-94: *Eucecryphalus* aff*. cervus*. (**G1**) Osh27: *Ceratocyrtis* cf*. galea*. (**G2**) Osh50: *Pseudodictyophimus clevei*. (**G3**) Vil269: *Acrobotrissa cribosa*. (**G4**) Mge17-134: *Lipmanella dictyoceras*. (**G5**) Ses60: *Peromelissa phalacra*. (**G6**) Vil512: *Pseudocubus obeliscus*. (**H1**) Osh129: *Cycladophora cornuta*. (**H2**) Mge17-6: *Valkyria*? *aurita*. (**I1**) Osh162: *Lamprotripus hirundo*. (**I2**) Vil225: *Pterocanium praetextum*. (**J1**) Vil444: *Theocorythium dianae*. (**J2**) Vil312: *Pterocorys carinata*.

The expertise required to collect, sort and identify living nassellarian specimens along with their short maintenance time in cultures (Anderson et al., 1989; Suzuki and Not, 2015) make the study of their taxonomy and ecology arduous. In addition, the low DNA concentration per individual cell challenges the molecular approach to address such questions. Phylogenetic studies have demonstrated the effectiveness of combining single cell DNA sequencing and imaging data in assessing classification and evolutionary issues beyond morphological characteristics (Bachy et al., 2012; Decelle et al., 2012; Biard et al., 2015). Acquisition of reference DNA barcode based on single cell sequencing of isolated specimens have also established the basis for further molecular ecology surveys inferring the actual diversity and ecology in the nowadays oceans (Decelle et al., 2013; Biard et al., 2016; Nitsche et al., 2016). To date, few phylogenetic studies have explored the extant genetic diversity of Nassellaria and the relationships among families and their evolutionary patterns remains still elusive (Kunitomo et al., 2006; Yuasa et al., 2009; Krabberød et al., 2011). So far, with a total of 16 sequences from morphologically described specimens, covering 5 of the 7 super-families identified, Eucyrtidioidea was considered as the most basal and the rest of the represented groups have uncertain phylogenetic positions (Krabberød et al., 2011).

The latest classification of Nassellaria (Matsuzaki et al., 2015) attempted to integrate the extensive morphological knowledge (Hertwig, 1879; Haeckel, 1882; Campbell, 1954; Petrushevskaya, 1971a; De Wever et al., 2001) with the few molecular analyses performed for Nassellaria (Krabberød et al., 2011). Here we introduce an integrative morpho-molecular classification of Nassellaria obtained by ribosomal DNA taxonomic marker genes (18S and partial 28S) and imaging techniques (Light microscopy, Scanning Electron Microscopy and/or Confocal Microscopy), compared with the current morphological classification. In addition, the extensive fossil record available for Nassellaria allowed us to time-calibrate our phylogenetic analysis based on molecular dating and infer their evolutionary history contextualized with geological environmental changes at a global scale. Finally, the inclusion of environmental sequences gave insights in the extant genetic diversity of nassellarians in contemporary oceans.

## Material and Methods

### Sampling and single cell isolation

Plankton samples were collected off Sendai (38° 0’ 28.8’’ N, 142° 0’ 28.8’’ E), Sesoko (26° 48’ 43.2’’ N, 73° 58’ 58.8’’ E), the Southwest Islands, South of Japan (28° 14’ 49.2’’ N, 129° 5’ 27.6’’ E), in the bay of Villefranche-sur-Mer (43° 40’ 51.6’’ N, 7° 19’ 40.8’’ E) and in the west Mediterranean (MOOSE-GE cruise) by net tows, Vertical Multiple-opening Plankton Sampler (VMPS) or Bongo net (20-300 μm). More information on sampling methodology can be found in the RENKAN database (http://abims.sb-roscoff.fr/renkan). Targeted specimens were individually handpicked with Pasteur pipettes from the samples and transferred 3-4 times into 0.2 μm filtered seawater to allow self-cleaning from debris, particles attached to the cell or preys digestion. Images of live specimens were taken under an inverted microscope and thereafter transfer into 1.5 ml Eppendorf tubes containing 50 μl of molecular grade absolute ethanol and stored at −20 ºC until DNA extraction.

### Single cell morphological identification

Nassellaria specimens were identified at the species level, referring to pictures of holotypes and reliable specimens, through observation of live images and posterior analysis of the skeleton by scanning electron microscopy and/or confocal microscopy. At the species level, we respected the following sentence of the International Code of Zoological Nomenclature 2000 (hereinafter ICZN 2000; http://www.iczn.org/iczn/index.jsp):

> the device of name-bearing types allows names to be applied to taxa without infringing upon taxonomic judgement […] For species and subspecies this name-bearing type is a single specimen […] for genera and subgenera it is a nominal species.

To determine the taxonomic position at the genus and family levels we followed the rules Articles 66 to 70 of “*types in the genus group*” in ICZN 2000: “*A nominal species is only eligible to be fixed as the type species of a nominal genus or subgenus*.” The family level was decided by the similarity of the type genus of the family based on ICZN 2000. This procedure has the advantage to directly use the fossil taxonomy (Matsuzaki et al., 2015) and the taxonomic concept of superfamily (Matsuzaki et al., 2015; Suzuki and Not, 2015) without ambiguity in the names used. Based on this rule, we updated the species name of specimens illustrated in previous studies and that are included in our phylogeny.

### DNA extraction, amplification and sequencing

DNA was extracted using the MasterPure Complete DNA and RNA Purification Kit (Epicentre) following manufacturer’s instructions. Once DNA was extracted and recovered, waste (i.e. pellet debris) from the extraction procedure were diluted in milliQ water to preserve skeletons and stored at −20°C. Both 18S rDNA and partial 28S rDNA genes (D1 and D2 regions) were amplified by Polymerase Chain Reaction (PCR) using Radiolaria and Nassellaria specific and general primers (Table 1). For further details about rDNA amplification see: http://dx.doi.org/10.17504/protocols.io.t5req56. PCR amplicons were visualized on 1% agarose gel stained with ethidium bromide. Positive reactions were purified using the Nucleospin Gel and PCR Clean up kit (Macherey Nagel), following manufacturer’s instructions and sent to Macrogen Europe for sequencing.

**Table 1.**
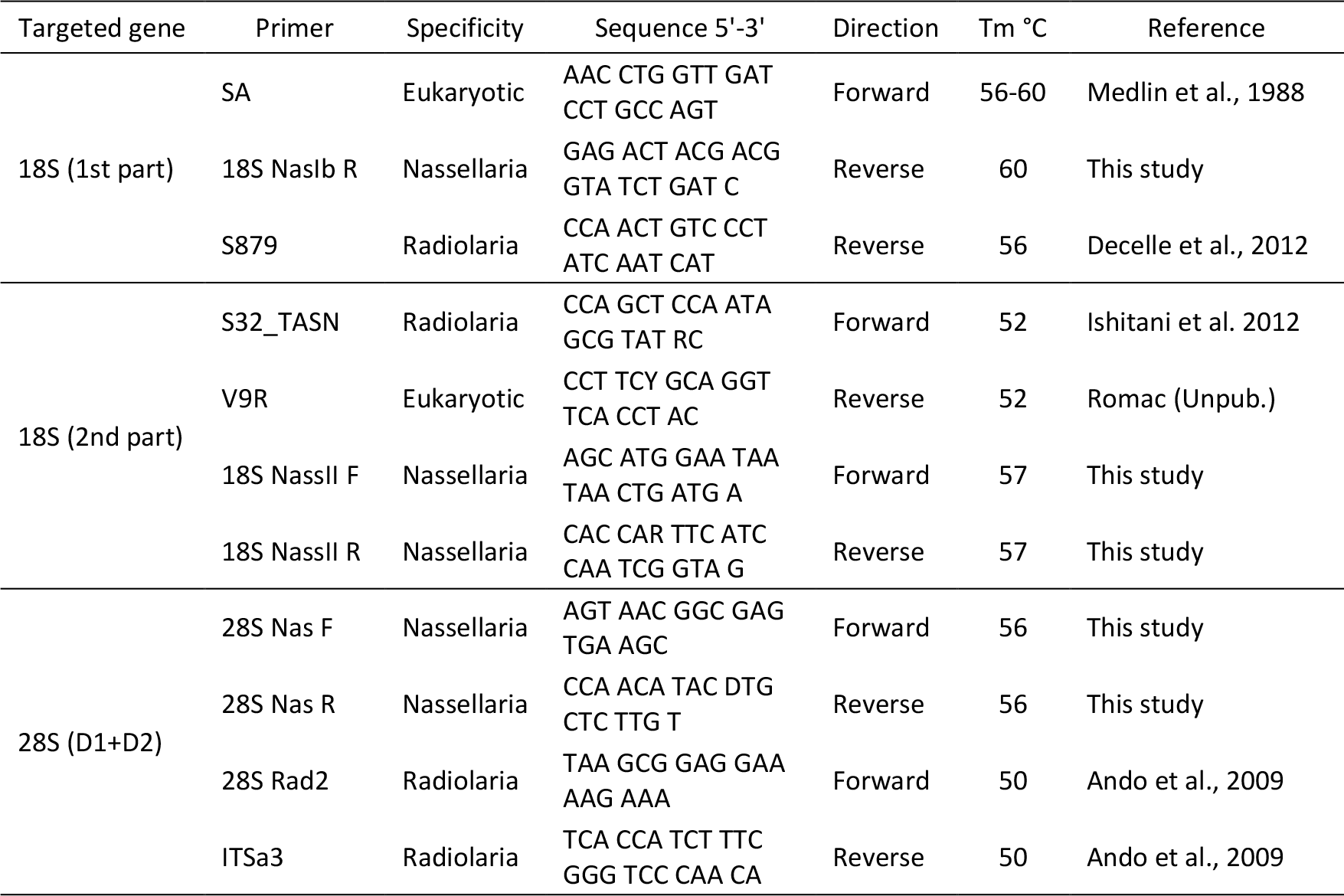
Primer sequences used for DNA amplification and sequencing.

### Phylogenetic analyses

After sequencing, forward and reverse sequences were checked and assembled using ChromasPro software version 2.1.4 (2017). Sequences were compared to reference database (GenBank) using the *BLAST search* tool integrated in ChromasPro to discriminate radiolarian sequences from possible contamination. Similar sequences identified in GenBank were retrieved and integrated in our databases. Two different datasets for each genetic marker (18S rDNA gene and partial 28S rDNA gene) were obtained and align separately using Muscle (Edgar, 2004) implemented in SeaView version 4.6.1 (Gouy et al., 2010) and manually checked. For both genes, the 18S rDNA (61 taxa, 1890 positions) and the 28S rDNA (57 taxa, 700 positions), phylogenetic analyses were performed independently. The best nucleotide substitution model was chosen following the corrected Akaike Information Criterion (AICc) using the *modelTest* function implemented in the R (R Core Team, 2014) package *phangorn* (Schliep, 2011). The obtained model (General Time Reversible with Gama distribution and proportion of Invariable sites, GTR+G+I) was applied to each data set in R upon the packages *APE* (Paradis et al., 2004) and *phangorn* (Schliep, 2011). A Maximun Likelihood (ML) method (Felsenstein, 1981) with 1000 replicates of bootstrap (Felsenstein, 1985) was performed to infer phylogenies. Since the topology and bootstrap support within main clades of both markers agree, the 18S rDNA and the 28S rDNA were concatenated in order to improve phylogenetic resolution. A final data set containing 90 taxa and 2590 positions was used to infer phylogenies following the previous methodology. The best model obtained was GTR+G+I, with 4 intervals of the discrete gamma distribution, and a ML method with 100 000 bootstraps were performed. In parallel, a Bayesian analysis was performed using BEAST version 1.8.4 (Drummond et al., 2012) with the same model parameters over 10 million generations sampled every 1000 states, to check the consistency of the topology and to calculate posterior probabilities. Final tree was visualized and edited with FigTree version 1.4.3 (Rambaut 2016). All sequences obtained in this study and used for the phylogeny were submitted in GenBank under the accession numbers: XXXXXXXX - XXXXXXXX.

### Molecular clock analyses

The resulting concatenated dataset of the 18S rDNA and the partial 28S rDNA used to infer phylogenies, was used for the molecular clock analyses. A Bayesian uncorrelated relaxed clock method (Drummond et al., 2006) implemented in BEAST version 1.8.4 (Drummond et al., 2012) was performed to calculate the divergence times of Nassellaria with a relaxed lognormal distribution. GTR+G+I with estimated base frequencies and 4 gamma categories was chosen for the substitution model, according to previous analyses (see section above). Both Spumellaria (outgroup) and Nassellaria nodes were forced to be monophyletic according to bootstrap results in the phylogenetic analysis. According to software options, a speciation Yule process (Yule, 1925; Gernhard, 2008) with random starting tree was chosen as tree prior, since our analyses are beyond population level. Markov chains were run in three parallel replicates for 100 million generations sampled every 1000 states and operators were left as default. Different replicates were combined in LogCombiner version 1.8.4 (Drummond et al., 2012) after removing the first 25% states. Twelve nodes were chosen to carry out this calibration (explained below from the oldest to the newest calibration age, and given the name of the node for the taxa they cover):

1. Root: The calibration for the root of the tree corresponds to the hypothesized last common ancestor between Nassellaria and Spumellaria (De Wever et al., 2001). A uniform distribution with a minimum bound of 200 million years ago (Ma) and a maximum of 600 Ma (U [200, 600]) was set to allow uncertainty in the diversification of both groups and to establish a threshold restricting the range of solutions for the entire tree.
2. Spumellaria: The second node corresponds to the origin of Spumellaria, the outgroup of our phylogenetic analyses. The first spumellarians appear in the fossil record in the Lower Ordovician (ca. 477.7 − 470 Ma) with the genera *Antygopora* (Aitchison et al., 2017). Therefore, the node was normally distributed with a mean of 474 and a standard deviation of 50 to avoid uncertainty due to the low genetic diversity of the outgroup: N (474, 50).
3. Nassellaria: The first nassellarian-like fossil appear in the Devonian (ca. 419.2 – 358.9 Ma) with internal spicules typical from Nassellaria (Cheng, 1986; Kiessling and Tragelehn, 1994; Schwartzapfel and Holdsworth, 1996). But it is not until the Early Triassic (ca. 250 – 240 Ma) that the first multi-segmented nassellarians appeared in the fossil record (Sugiyama, 1992, 1997; De Wever et al., 2003; Hori et al., 2011; Dumitrica, 2017). Because it is unclear whether the first nassellarian-like fossils are true nassellarians, a uniform distribution with a maximum boundary of 500 and a minimum of 180 (U [180, 500]) was chosen to allow uncertainty in the origin of living Nassellaria (Suzuki and Oba, 2015).
4. Eucyrtidioidea N (246, 10): The first appearance of this family in the fossil record is with the genus *Triassocampe* (Sugiyama, 1997) in the early Middle Triassic (Anisian: ca. 246.8 − 241.5). And there are continuous fossil record since the Carnian (ca. 237 – 228.5 Ma; as well Late Triassic) to present of the genus *Eucyrtidium* (Petrushevskaya, 1971a; De Wever et al., 2001, 2003). Hence, the node was normally distributed with an average mean of 246 and a standard deviation of 10 (N (246, 10)).
5. Lophophaenidae N (191, 10): This family belongs to the Superfamily Plagiacanthoidea, and the genus *Thetisolata* in the Early Jurassic (Pliensbachian: ca. 191.4 – 183.7 Ma) is the oldest fossil record associated to this family (De Wever, 1982).
6. Artostrobioidea N (182, 10): The genus *Artostrobium*, questionably assigned by Matsuoka (2004), is the oldest fossil for this family, and its first appearance in the fossil record is dated in the Early Jurassic (Toarcian: ca. 183.7 − 174.2 Ma) (Matsuoka, 2004).
7. Sethoperidae N (180, 30): Like Lophophaenidae, this family also belongs to the Plagiacanthoidea, but its first appearance in the fossil record dates in the Middle Jurassic (Bajocian: ca. 170.3 – 168.3 Ma) with the genus *Turriseiffelus* (Dumitrica and Zügel, 2003). However, the fossil record of this family seems to be patched, since the very first member associated to this family is restricted to a single stage in the Early Jurassic (genus *Caterwhalenia*, Pliensbachian: ca. 191.4 – 183.7) (O’Dogherty and Carter, 2009). Due to this uncertainty in the appearance of this family, we decided to increase the node prior standard deviation to 30 Ma.
8. Cannobotryoidea N (122, 20): Petrushevskaya (1971b) dated the first appearance of this family in the Cretaceous (ca. 145 – 66 Ma), and later the time frame was restricted to the Early Cretaceous (ca. 145 – 100 Ma) (De Wever et al., 2001, 2003). However, there is no consensus about whether the first morphologically similar genera (*Ectonocorys* or *Solenotryma*) to this group that appear in the fossil record really belong to it or not. Therefore, since no adequate evidence has been shown we slightly increase the standard deviation (sd: 20 Ma) for this node prior to allow uncertainty (O’Dogherty et al., 2009; Matsuzaki et al., 2015).
9. Acanthodesmoidea N (66, 10): The three families belonging to this superfamily (Acanthodesmiidae, Stephaniidae, Triospyrididae) have their first appearance in the fossil record in the Paleocene (ca. 66 − 56) (Petrushevskaya, 1971a; De Wever et al., 2001, 2003), and the first genus described associated to this families corresponds to *Tholospyris* in the Early Paleocene (Kozlova, 1983, 1999).
10. Bekomidae N (62, 10): The genus *Bekoma* (as representative of the family Bekomidae) has its first appearance in the Late Paleocene (ca. 66 − 56) (Nishimura, 1992; De Wever et al., 2001).
11. Pterocorythidae N (57, 10): *Cryptocarpium* is the first, but doubted, representative of this family in the fossil record and it is dated in the Late Paleocene (ca. 66 − 56 Ma), followed by *Podocyrtis*, the first true representative, in the transition of the Eocene (ca. 56 – 33 Ma) (Sanfilippo and Riedel, 1992; De Wever et al., 2001; Hollis, 2006).
12. Carpocaniidae N (38, 10): The first representative of this family appears in the late Eocene (ca. 38 – 33.9 Ma) and correspond to the genus *Carpocanium* (De Wever et al., 2001; Kamikuri et al., 2012).

### Post hoc analyses

In order to provide statistical support to our conclusions two different analyses were developed: a diversification of taxa over time (Lineages Through Time: LTT) and an ancestral state reconstruction. The former analysis was carried out with the *ltt.plot* function implemented in the package *APE* (Paradis et al., 2004) upon the tree obtained by the molecular dating analyses after removing the outgroup. The second analysis uses the resulting phylogenetic tree to infer the evolution of morphological characters. A numerical value was assigned to each state of a character trait, being 0 for the outgroup or the considered ancestral state, and 1 to 4 for Nassellaria or the presumed divergence state. In total 5 traits were consider (Table 2): the number of cyrtids/segments (1-monocyrtid, 2-dicyrtid, 3-tricyrtid, 4-multicyrtid), the complexity of the cephalis (1-simple / spherical, 2-hemispherical / elongated / reduced / fussed with thorax, 3-Lobes, 4-complex), the apical (A) ray, the ventral (V) ray (1-not projecting the skeleton, 2-small, 3-medium/big, 4-united with A/V) and the dorsal (D) and lateral rays (Lr and Ll) treated as one single character (D+Lr+Ll: 1-not projecting the skeleton, 2-small, 3-medium, 4-big/foot/wings). Once the character matrix was established a parsimony ancestral state reconstruction was performed to every character independently.

**Table 2.** List of morphological characters (traits) and their states (1 to 4) used for the ancestral state reconstruction analysis.

|  |  | Traits |  |  |  |  |
| --- | --- | --- | --- | --- | --- | --- |
|  |  | Number of segments | Complexity of cephalis | Apical ray | Ventral ray | Dorsal and lateral rays |
| State | 1 | Monocyrtid | Simple, Spherical | Not projecting the skeleton | Not projecting the skeleton | Not projecting the skeleton |
|  | 2 | Dicyrtid | Hemispherical, elongated, Reduced, fused with thorax | Small | Small | Small |
|  | 3 | Tricyrtid | Lobes | Medium, big | Medium, big | Medium |
|  | 4 | Multicyrtid | Complex | United with V | United with A | Big, foot, wings |

### Environmental sequences

Each nassellarian sequence of the 18S rDNA and partial 28S rDNA was compared with publicly available environmental sequences in GenBank (NCBI) using BLAST (as of December 2017); in order to estimate the environmental genetic diversity of Nassellaria and eventually to check the genetic coverage of our phylogenetic tree. Environmental sequences were placed in our reference phylogenetic tree using the pplacer software (Matsen et al., 2010). A RAxML (GTR+G+I) tree was built for the placement of the sequences with a rapid bootstrap analysis and search for best-scoring ML tree and 1000 bootstraps.

### Confocal (CM) and Scanning Electron Microscopy (SEM)

After DNA extraction, nassellarian skeletons were recovered from the eluted pellet and handpicked under binoculars or inverted microscope. After cleaning and preparing the skeletons (detailed protocol in http://dx.doi.org/10.17504/protocols.io.t5meq46 for SEM and http://dx.doi.org/10.17504/protocols.io.t5qeq5w for CM) images were taken with an inverted Confocal Microscopy (Leica TCS SP5 AOBS) and/or FEI Phenom table-top Scanning Electron Microscope (FEI technologies).

## Results

### Comparative molecular phylogeny and morphological taxonomy

A total of 90 distinct nassellarian specimens generated 61 sequences for the 18S rDNA gene and of 57 sequences for the partial 28S (D1 & D2 regions) rDNA gene (Supplementary Material Table S1). The alignment matrix has 32.4% of invariant sites, and includes 67 new sequences and 23 reference sequences, of which 7 are environmental and 16 have been morphologically identified. All of these specimens cover 7 superfamilies (Acanthodesmoidea, Acropyramioidea, Artostrobioidea, Cannobotryoidea, Eucyrtidioidea, Plagiacanthoidea, Pterocorythoidea, and three undefined families), based on morphological observations performed with light microscopy (LM), scanning electron microscopy (SEM) and/or confocal microscopy (CM) on the exact same specimens for which we obtain a sequence. Overall, molecular phylogeny is consistent with morphological classification at the superfamily level, although there are some specific discrepancies. The phylogenetic analysis shows 11 clades (Fig. 1) clearly differentiated with high values of ML bootstrap (BS > 99) and posterior probabilities (PP > 0.86).

Clade A holds the most basal position with 16 sequences of which 10 are novel sequences, 5 were previously available and one is environmental. All specimens clustering within this clade (Fig. 2.A, Fig. 3.A) have a simple and round cephalis, an apical horn, a small ventral rod and multisegmented (cephalis, thorax and several abdomen) skeleton with distinctive inner rings, that correspond to the Superfamily Eucyrtidioidea. All multisegmented nassellarians with spherical cephalis encountered in our study belong to this superfamily, both morphologically and phylogenetically. The rest of the clades group together with a high BS (100) and PP (1) values. Thereafter clades B, C, D and E cluster together in lineage II. Clades X, F and G constitute the lineage III, highly supported (100 BS and 1 PP) as sister group of the lineage IV. This last lineage it is composed by the clades H, as the basal group and clades I and J highly related phylogenetically (100 BS and 1 PP).

**Figure 3.**
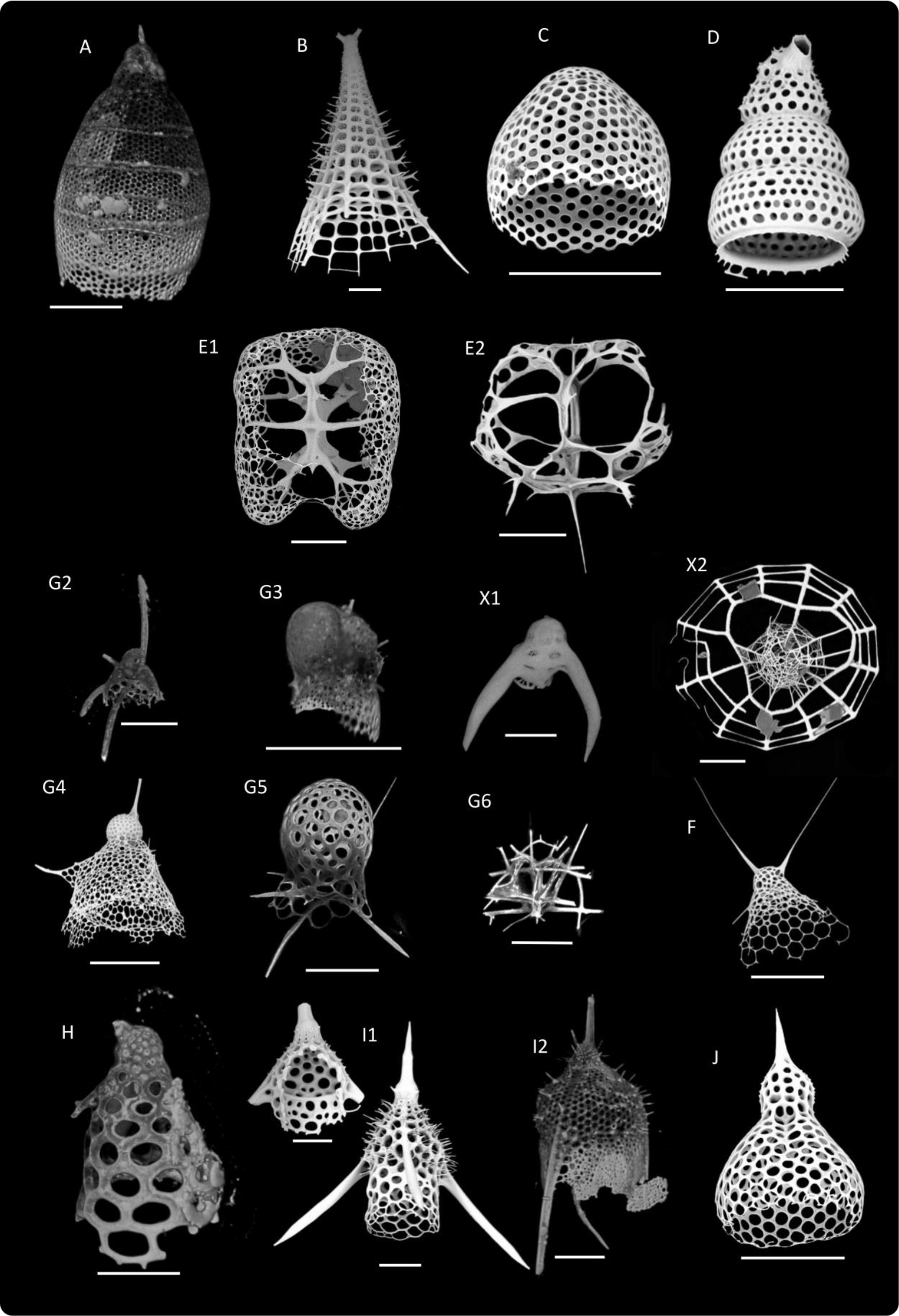
Scanning Electron Microscopy (SEM) and/or Confocal Microscopy (CM) images of Nassellarian specimens used in this study for phylogenetic analysis or morphologically related to one of the morpho-molecular clades of Fig. 1. Scale bars = 50μm. (**A**) Vil496: *Eucyritidium acuminatum* (CM). (**B**) Osh128: *Polypleuris giganthea* (SEM). (**C**) Mge17-122: *Carpocanium* cf. *obliqua* (SEM). (**D**) Mge17-70: *Botryostrobus auritus* (SEM). (**E1**) Toy231: *Amphispyris costata* (SEM). (**E2**) Vil267: *Lophospyris* aff*. ivanovi* (SEM). (**X1**) Osh112: *Nothotripodiscinus johannismonicae* (CM). (**X2**) Osh130: *Enneaphormis enneastra* (SEM). (**F**) Mge17-94: *Eucecryphalus* aff. *cervus* (SEM). (**G2**) Osh50: *Pseudodictyophimus clevei* (CM). (**G3**) Vil269: *Acrobotrissa cribosa* (CM). (**G4**) Mge17-137: *Lipmanella* sp. (SEM). (**G5**) Ses60: *Peromelissa phalacra* (CM). (**G6**) Vil512: *Pseudocubus obeliscus* (CM). (**H**) Osh129: *Cycladophora cornuta* (CM). (**I1 left**) Osh184: *Lamprotripus hirundo* (SEM). (**I1 right**) Osh127: *Lamprotripus hirundo* (SEM). (**I2**) Vil225: *Pterocanium praetextum* (CM). (**J1**) Vil444: *Theocorythium dianae* (SEM).

Within lineage II, clade B is represented by only one sequence from this study (Osh128), and its morphology matches with the superfamily Acropyramioidea (Fig. 2.B, Fig. 3.B) exhibiting a pyramidal skeleton constituted of a reduced cephalis and thorax. Clade C is composed by two novel sequences (Fig. 2.C, Fig. 3.C). Their overall round morphology and a characteristic small and flat cephalis not well distinguished from the thorax agrees with the undetermined family Carpocaniidae (Petrushevskaya, 1971a; De Wever et al., 2001). Its sister clade, the Clade D is constituted by three new (Ses58, Mge17-70 and Mge17-9) and one environmental sequences, and the morphology of these specimens (Fig. 2.D, Fig. 3.D) agrees with the definition of the Superfamily Artostrobioidea.

They share a multisegmented skeleton without significant inner rings, a hemispherical cephalis and an important ventral rod. The last clade of the lineage II, clade E, gathers four new sequences, one and two environmental sequences. This clade includes all the monocyrtid (cephalis) nassellarians where the spines A and V are merged forming the so-called D-ring (Acanthodesmoidea). There are representatives for two out of three families (Stephaniidae is the missing family), yet no phylogenetic differences were found for the included families, Acanthodesmiidae (Fig. 2.E1, Fig. 3.E1) and Triospyrididae (Fig. 2.E2-E4, Fig. 3.E2).

In lineage III, clades X and F are highly supported as sister clades (100 BS and 1 PP). Clade X is established by two novel sequences morphologically identified as *Nothotripodiscinus johannismonicae* (Fig. 2.X1, Fig. 3.X1) and *Enneaphormis enneastra* (Fig. 2.X2, Fig. 3.X2). These two specimens have a very short or missing median bar (MB) allowing the development of three large feet with a three-pointed star (dorsal and lateral, left and right, rays) shape forming a significant circular frame where they build the thorax when present. In Clade F there are two novel sequences (Ses59 and Mge17-94) and a previously available sequence. These specimens are morphologically identified in the genus *Eucecryphalus* (Fig. 2.F, Fig. 3.F) and a third specimen as *Tricolocamptra* sp. (AB246685). The former genus belongs to the undetermined family Theopiliidae (De Wever et al., 2001; Matsuzaki et al., 2015) with a hat shaped morphology and pronounced segmentation. The genus *Eucecryphalus*, the senior synonym of *Theopilium*, is the type genus of the family Theopiliidae (Matsuzaki et al., 2015), and thus Clade F is automatically specified as the “true” Theopiliidae. The taxonomic position of the second genus, *Tricolocamptra*, varies through references and the available image for the specimen lacks taxonomical resolution, therefore it is not possible to establish the link with any described morphological group.

With 14 different morpho-species and 12 different genus, Clade G is the most morphologically diverse clade. While all specimens have consistently one or two segments, the complexity of the cephalis and the sizes of the rays vary widely (Fig. 2.G1-G6, Fig. 3.G2-G6). Specimen HQ651793 (*Archiscenium tricolpium*) corresponds to the Family Sethophormididae (Sethophormididae Haeckel, 1882, *sensu emend*. Petrushevskaya, 1971a) within the Superfamily Plagiacanthoidea. The large cephalis and the umbrella shape of the thorax are characteristics of this family. The specimen Vil269 (Fig. 2.G3, Fig. 3.G3, *Acrobotrissa cribosa*) agrees with the description of the Superfamily Cannobotryoidea and its complex cephalic structure subdivided in different lobes is the main characteristic. The rest of the specimens of clade G can be grouped within the Family Lophophaenidae (Lophophaenidae Haeckel, 1882, *sensu* Petrushevskaya, 1971a), still within Plagiacanthoidea. Within this family there are two different subclades, those specimens with a prominent thorax (e.g. Mge17-134, *Lipmanella dictyoceras*, Fig. 2.G4, Mge17-137, *Lipmanella* sp. Fig. 3.G4) and those with a barely defined thorax (e.g. Ses60, *Peromelissa phalacra*, Fig. 2.G5, Fig. 3.G5) or not-defined thorax (e.g. Vil512, *Pseudocubus obeliscus*, Fig. 2.G6, Fig. 3.G6). Two last specimens, Osh27 (*Ceratocyrtis* cf. *galea*, Fig. 2.G1) and Osh50 (*Pseudodictyophimus clevei*; Fig. 2.G2, Fig. 3.G2), are morphologically identified as Lophophaenidae (HQ651793: De Wever et al., 2001; & Osh50: Matsuzaki et al., 2015; Petrushevskaya, 1971b) despite their phylogenetic position closely related to the Superfamily Cannobotryoidea.

Lineage IV is the most distal regarding the root of the tree. Within it, clade H is constituted by six new sequences very closely related to each other (BS>96 and PP>0.99). These specimens are morphologically assigned to the genus *Cycladophora* (Fig. 2.H1, Fig. 3.H) and *Valkyria* (Fig. 2.H2). Specimens from clade H show a conical and campanulate shaped morphology with a distinguishable cephalis and an apical horn compared to the wide thorax and not distinguished cephalis with two (V and A) rods of *Eucecryphalus*. Clade I is composed of six novel sequences and one previously available, while clade J gathers twenty new sequences and 3 previously available. These clades comprise nassellarians with an apical stout horn, a spherical (clade I) or elongated cephalis (clade J) and a truncated conical thorax; the abdomen, if present, is also truncated and sometimes not well defined, agreeing with the definition of the Superfamily Pterocorythoidea. Clade J corresponds to those with elongated cephalis (Fig. 2.J, Fig. 3.J) or Pterocorythidae (Pterocorythidae Haeckel, 1882, *sensu* De Wever et al., 2001). Clade I includes nassellarians with spherical head, ventral short ray and three feet (Fig. 2.I2, Fig. 3.I2), all characteristics of the Family Lychnocanidae (Lychnocaniidae Haeckel, 1882, *sensu* emend. Suzuki in Matsuzaki et al. 2015). Two specimens (Fig. 2.I1, Fig. 3.I1) of the undetermined family Bekomidae (Bekomidae Dumitrica in De Wever et al., 2001), are scattered among clade I showing no phylogenetic differences between members of both families. The BS (93) and the PP (0.99) values establish a strong phylogenetic relationship within this lineage, and so does the overall skeleton shape.

### Molecular dating

The molecular clock dated the diversification between Spumellaria and Nassellaria (the root of the tree) with a median value of 512 Ma (95% Highest Posterior Density -HPD-: between 600 and 426 Ma) (Fig. 4). From now on, all dates are expressed as median values followed by the 95% HPD interval. Despite the large uniform distribution given to the ingroup (U [180, 500]), the first diversification of Nassellaria was settled at 423 (500-342) Ma. Clade A had its first radiation dated in 245 (264-225) Ma. The common ancestor to all the other clades diversified at around 340 (419-271) Ma into two main groups, the so-called lineage II and the lineages III-IV splitting into two other lineages soon afterwards. Within lineage II, the first diversification occurs at 276 (354-209) Ma, and clades C and D diversified 197 (250-160) Ma. Thereafter in this lineage the phylogenetic relationships are dubious, however clade D diversified from any other clade 248 (315-191) Ma. Diversification within these clades was 28 (49-4) Ma for clade C, 175 (194-155) Ma for clade D and 77 (95-60) Ma for clade E. The lineages III and IV diversified from each other 274 (344-215) Ma. Lineage III rapidly diversified 243 (304-197) Ma when clades F and X split from clade G, followed by the fast diversification of clade G 196 (241-170) Ma. It was 168 (243-76) Ma when clade F separate of clade X, and 86 (157-32) Ma and 40 (106-4) Ma when they respectively diversified. The last lineage (IV) diversified between clades H, I and J 206 (287-129) Ma, and clades I and J 138 (206-88) Ma. Despite this early divergence between clades, radiation within clades was more recent, being 87 (171-26) Ma, 70 (87-53) Ma and 73 (90-58) Ma for clades H, I and J, respectively.

**Figure 4.**
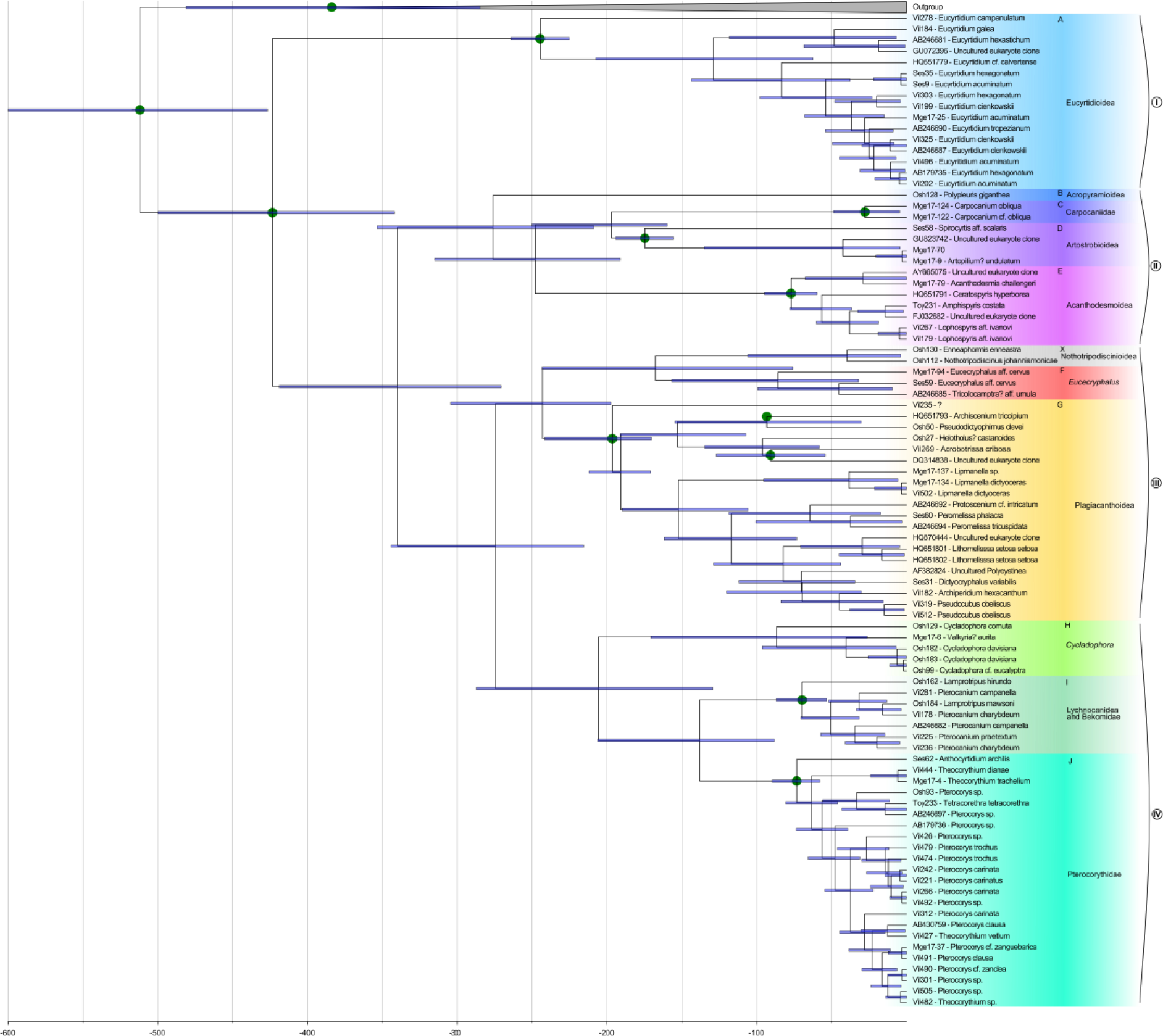
Time-calibrated tree (Molecular clock) of Nassellaria, based on alignment matrix used for phylogenetic analyses. Node divergences were estimated with a Bayesian relaxed clock model and the GTR + γ + I evolutionary model, implemented in the software package BEAST. Twelve different nodes were selected for the calibration (green dots). Blue bars indicate the 95% highest posterior density (HPD) intervals of the posterior probability distribution of node ages.

### Post-hoc analyses

The lineages through time analysis (Fig. 5) shows a classic exponential diversification slope, with a 0.0114 rate of speciation (Ln(lineages)·million years^−1^). The first diversification of extant Nassellaria (~394 Ma) corresponds to the divergence between clade A and the rest of the clades. However, the first increase in the slope (up to: 0.017, t-test: p<0.001) occurs at ~265 Ma, when the main lineages start expanding; lineage II splits between clade B and clades C, D and E, clade A diversifies and the evolutionary lineages III and IV diverged from each other followed by the rapid diversification of lineage III. After this sudden increase of the main lineages, a second diversification event happened (ca. 198 Ma) where both the evolutionary lineage IV and the clade G diversify and clade C splits from clade D. Thereafter, the diversification seems to be stepped and separate in different periods of time, were only lineage III keep diversifying. The last and relatively uninterrupted diversification occurred ~82 Ma corresponding to the speciation within the already present clades and the first diversification of the clades H, F, E, J, I, X and C.

**Figure 5.**
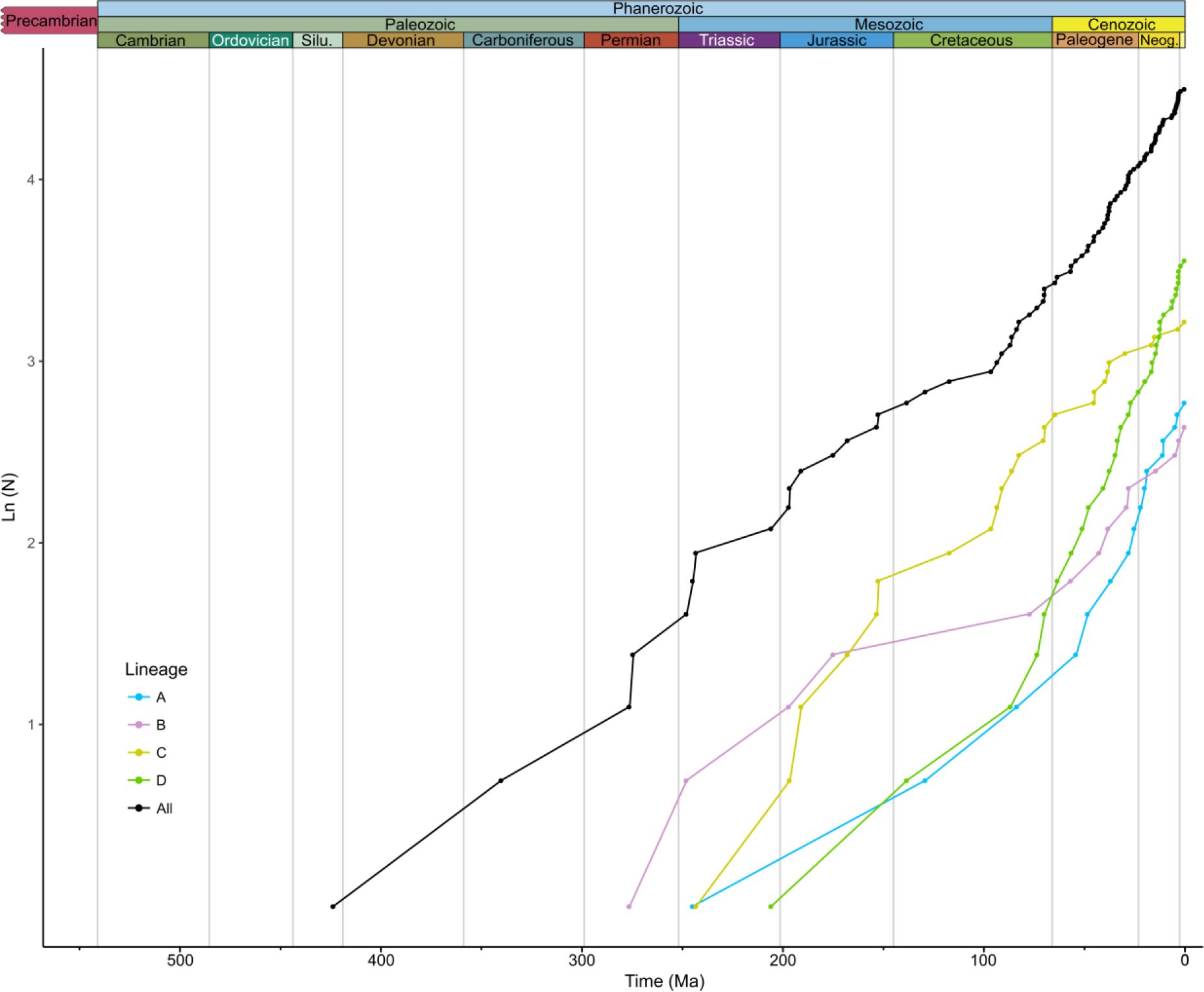
Lineages Through Time (LTT) analysis based on the molecular clock results for Nassellaria (removing the outgroup; in black), and of each lineage independently: lineage A (Blue), lineage B (purple), lineage C (yellow) and lineage D (green).

The ancestral state reconstruction analyses (Fig. 6) establish that the common ancestor to all living Nassellaria should be either multicyrtid or dicyrtid, with 1^st^ or 2^nd^ cephalic state (Table 2), a small apical horn, with a small or not projecting ventral ray and not projecting dorsal and lateral rays out of the skeleton. Soon after the diversification between clade A and the rest of the clades, the analysis establishes for the common ancestor of lineages II, III and IV, a dicyrtid state, with a 2^nd^ cephalic state (Table 2), a small apical horn, and not projecting ventral, dorsal and lateral rays. The same morphology was found for the common ancestor of the lineage II, for that of lineages III and IV and that of lineage III. The dicyrtid state remains the most parsimonious state for the common ancestor of all families except for clades A and D and E. The cephalis has a 2^nd^ state (Table 2) for every common ancestor of each clade except for clade A (1^st^ state; Table 2, F) (1^st^ or 2^nd^; Table 2) and I-J (2^nd^ or 3^rd^; Table 2). Regarding states of the apical ray, it remained small in every node with the exception of lineage IV, where it is medium/big. The ventral ray remained inside the skeleton with the exception of clade A, D, G and I, where it is small. Finally, the dorsal and lateral rays remained inside the skeleton for the common ancestor of clades A, B, C, D and F, but quite different states for the common ancestor of the rest of the clades: clade E has either a not projecting or small dorsal & lateral rays, clade X is characterized by big/feet/wings, clade G has a small dorsal and lateral rays, clades H and J either not projecting out of the skeleton, small or medium dorsal and lateral rays and clade I has a big/feet/wings dorsal and lateral rays.

**Figure 6.**
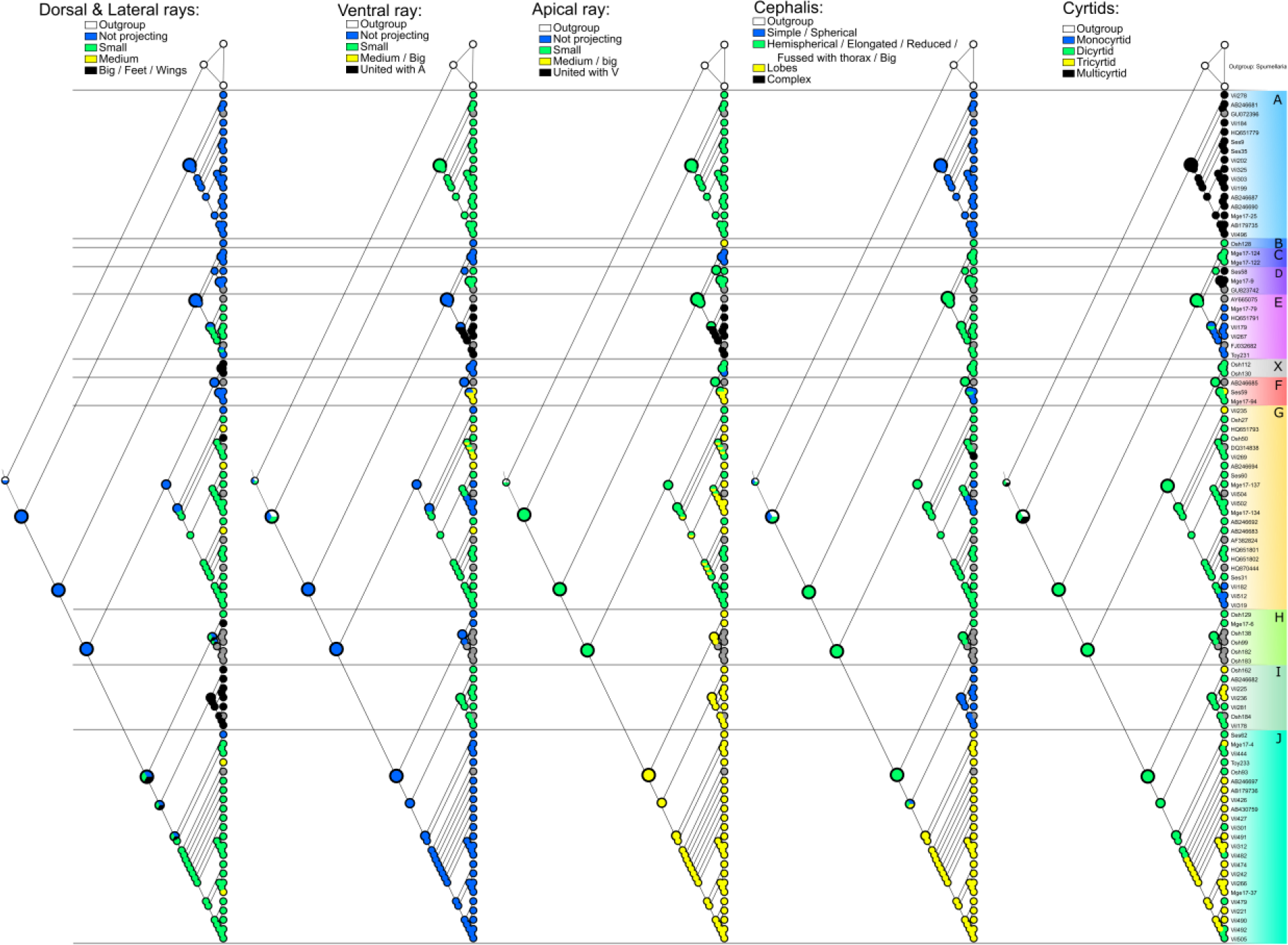
Parsimony Ancestral State Reconstruction analysis based in the resulting phylogenetic tree for the 5 characters chosen. Relevant nodes are increased for clarity.

### Environmental genetic diversity of Nassellaria

A total of 229 18S rDNA and two 28S rDNA environmental sequences affiliated to Nassellaria were retrieved from public database and placed in our reference phylogenetic tree (Fig. 7; Supplementary Material Table S2). Most of the environmental sequences (151) belonged to clade G, while 41 other sequences were scattered between clades A, D, E, F, I and J (2, 6, 26, 1, 2 and 4 sequences respectively). The rest of the sequences could not be placed within any existing clade. From those, 28 were closely related to clade E, and the others at basal nodes mainly over lineage II. These 37 sequences were then included in a phylogenetic tree (with a GTR+G+I model, 4 invariant sites and 1000 bootstraps) and 3 new and highly supported clades appeared (Supplementary Material Fig. S1). The first clade, constituted by 31 sequences, was assigned to Collodaria in Biard et al. (2015). A second clade formed by 2 sequences (BS: 94) was highly related to clades B and C (BS: 96). And the last clade, constituted as well of 2 sequences (BS: 100) was basal to the node formed by clades C, D and the previous mentioned environmental clade (BS: 58). The 2 remaining sequences were highly related to clades I (BS: 84) and to the node comprising clades I and J (BS: 100). Finally, 2 sequences were aligned within clade D and the last sequence was closely related to clade B, C and D. Regarding the sequences blasted for the partial 28S matrix, 2 sequences were extracted and mapped within clades E and G respectively.

**Figure 7.**
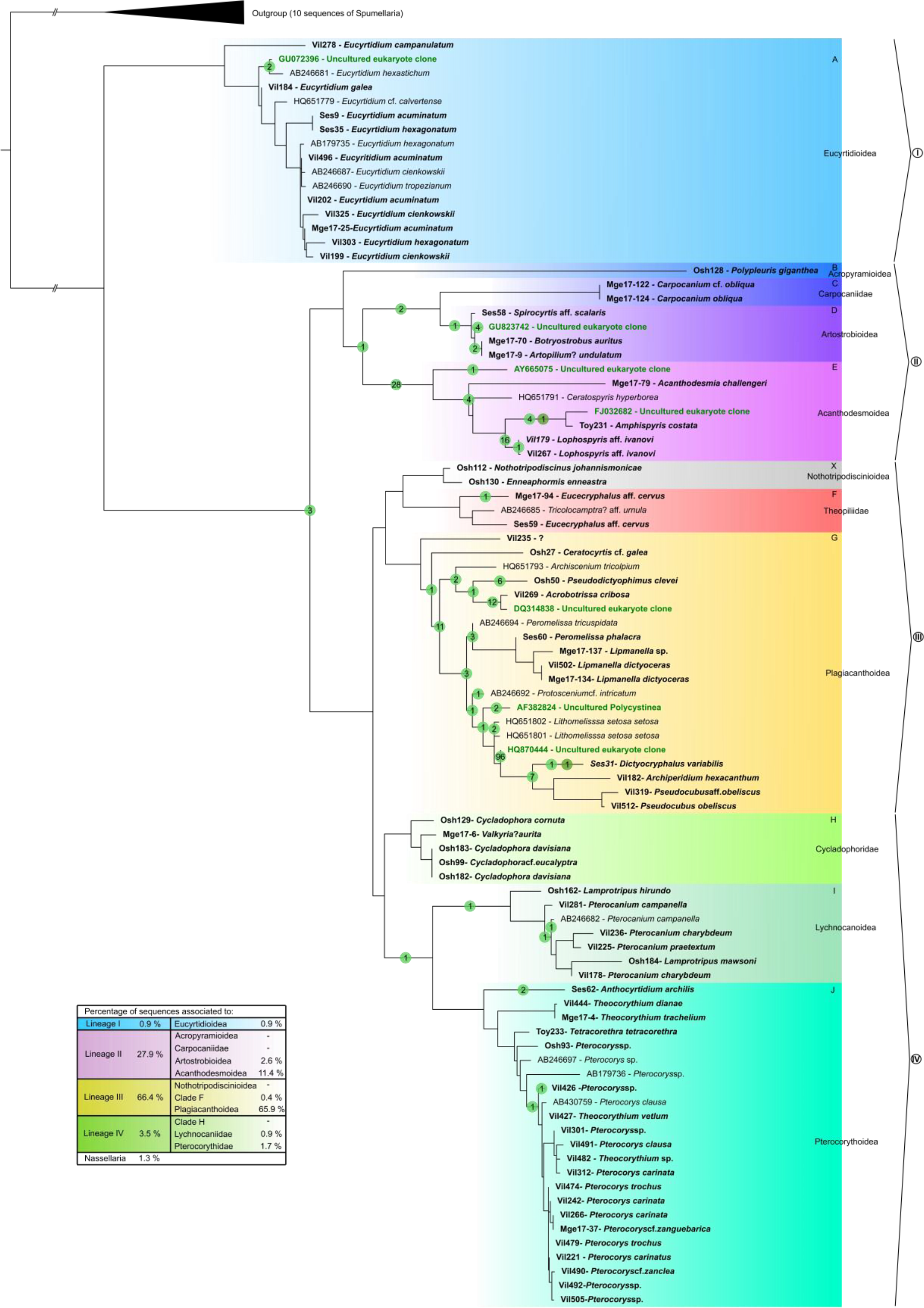
Pplacer phylogenetic placement of 229 environmental sequences for the 18S rDNA gene and 2 environmental sequences for the 28S rDNA gene into a concatenated phylogenetic tree of Nassellaria (complete 18S + partial 28S rDNA genes). Numbers at nodes represent the amount of environmental sequences assigned to a branch or a node; in light green for the 18S rDNA gene and in dark green for the 28S rDNA gene.

## Discussion

### Morpho-molecular classification of living Nassellaria

In our phylogenetic classification the overall morphology of Nassellaria, rather than the initial spicular system, is the most accurate feature to differentiate clades at the Superfamily level. We could not find any pattern in the initial spicular system that enables to separate clades, suggesting that the complexity of the initial spicular system is not related to Nassellaria phylogeny (e.g. Acanthodesmoidea, Plagiacanthoidea). Yet, this morphological character has been used to discriminate families and higher taxonomical levels (Petrushevskaya, 1971b). As well, the arches connecting these spicules were used in previous classifications and, in some cases, to discern at genus, and partly family levels (Petrushevskaya, 1971b; De Wever et al., 2001). In addition to our results, it has never been hypothesized the evolution of the initial spicular system complexity through the radiolarian fossil record. The use of the overall morphology in Nassellaria classification makes recognition of live cells easier under a light microscope facilitating morphology based ecological studies.

The extent Nassellaria included in our study can be divided in 11 morpho-molecular clades, grouped in four main evolutionary lineages based on phylogenetic clustering support, common morphological features and molecular dating: Eucyrtidioidea in lineage I; Acropyramioidea, Carpocaniidae, Artostrobioidea and Acanthodesmoidea in lineage II; Nothotripodiscinioidea, Theopiliidae and Plagiacanthoidea in lineage III, and *Cycladophora*, Lychnocanoidea and Pterocorythoidea in lineage

IV. The unified morpho-molecular framework here proposed reveals a partial agreement between the traditional taxonomy and the molecular classification. Such discrepancies have already been reported not only in other Radiolaria groups (Decelle et al., 2012; Biard et al., 2015) but as well in other SAR taxa such as Foraminifera (Pawlowski and Holzmann, 2002; Pawlowski et al., 2003), Phaeodaria (Nakamura et al., 2015) or Tintinnida (Bachy et al., 2012), being a common issue in protists classification (Schlegel and Meisterfeld, 2003; Caron, 2013).

Our revised morpho-molecular classification confirms the monophyly of the ancient Superfamilies Eucyrtidioidea and Acropyramioidea, represented by the clades A and B respectively, as well as the undetermined Family Carpocaniidae, the Superfamilies Artostrobioidea and Acanthodesmoidea, (clades C, D and E, respectively). The Superfamily Plagiacanthoidea is however paraphyletic, appearing in two different clades (X and G). Similarly, previously proposed families are scattered within clade G or even including the Superfamily Cannobotryoidea. Clade F and H both display an identical architecture of the initial spicular system, yet they are clustered based in the overall morphology in two different clades, the family here proposed as Theopiliidae (clade F) and *Cycladophora*-like specimens (clade H). As the genus *Eucecryphalus* (clade F) is the genus of the family Theopiliidae (see Matsuzaki et al., 2015), clade F holds the name Theopiliidae. Regarding clade H, a new family Cycladophoridae is established herein as defined in the taxonomic note.

Regarding clades I and J, Matsuzaki et al. (2015) include them within the same Superfamily (Pterocorythoidea) due to the strong phylogenetic relationship reported by Krabberød et al. (2011). Yet due to the evolutionary patterns and the phylogenetic distance, these two clades should be considered as two different superfamilies, Pterocorythoidea and Lychnocanoidea (Lychnocanoidea Haeckel, 1882, sensu Kozur and Mostler, 1984). In our study the undetermined family Bekomidae was scattered within clade I, showing no phylogenetic differences. Re-examination of the cephalic structure in *Lamprotripus* concludes the assignation of the genus to the Superfamily Lychnocanoidea, yet intergeneric morphological differences remain still elusive. Further molecular analyses must reveal these discrepancies between morphological and molecular classification. For example, differences in the previous clade can be due to an environmental differentiation and/or the effect of hosting photosynthetic symbionts; since all specimens belonging to the genus *Pterocanium* were collected in surface waters of Villefranche-sur-Mer (Mediterranean Sea), whereas those of *Lamprotripus* come from deep samples (2000-3000m) off Sendai (Japan).

### Evolutionary history of Nassellaria

The morphological evolution of Nassellaria is marked by their dubious appearance in the fossil record, whether it happened with the primitive nassellarian forms (in the Upper Devonian, ca. 372 Ma) or with the first multi-segmented nassellarians (in the Early Triassic, ca. 251.9 Ma) (De Wever et al., 2001; Suzuki and Oba, 2015). Our results showed that the first diversification of Nassellaria agrees in both time (~423 Ma; 95% HPD: 500-342 Ma; Figs. 4, 5) and reconstructed morphology (two- or multi-segmented last common ancestor) with the primitive nassellarians. Thus, the most likely scenario, proposed by Petrushevskaya (1971) and continued by Cheng (1986), is where Nassellaria originated during the Devonian from primitive radiolarian forms and not during the Triassic. This hypothesis can explain the sudden appearance of forms in the Middle Triassic (ca. 246.8 Ma) already pointed out by De Wever et al. (2003) and confirmed by our results.

Despite the early appearance of the Order Nassellaria, living nassellarian groups diversified during the Early Mesozoic (ca. 250 Ma) after the third, and biggest of all, mass extinction (Sepkoski, 1981; Bambach et al., 2004). This bottle neck led to a rapid increase in the global marine diversity (Twitchett et al., 2004), where all the surviving and isolated populations followed different pathways; an expected process in the aftermath of an evolutionary crisis (Twitchett and Barras, 2004; Hull, 2015). The lineage of Acropyramioidea/Acanthodesmoidea (lineage II) splits apart into many different forms that will led to new families, lineages III and IV diverge from each other followed by a fast branching of the plagiacanthoids, while eucyrtidioids (lineage I) slowly diversifies. By the end of the Late Triassic-Early Jurassic, radiolarian fossil record reaches its highest diversity measures (De Wever et al., 2006). Along with Nassellaria, other groups of marine protists also diversified such as dinoflagellates (Fensome et al., 1996; Janouškovec et al., 2017) or even appeared like diatoms (Kooistra and Medlin, 1996; Sims et al., 2006).

The second part of the Mesozoic is characterized by a global oceanic anoxic event happening at the end of the early Jurassic (ca. 183.7 − 174.2 Ma; Jenkyns, 1998) and a widespread series of Oceanic Anoxic Events (known as OAEs) during the Cretaceous (ca. 145-66 Ma; Jenkyns, 2010; Schalanger and Jenkyns, 1976). These events have been proposed as the major responsible force for the appearance of planktic Foraminifera during the Jurassic (Hart et al., 2003), and for extinction-speciation processes in planktonic evolution during the Cretaceous, especially for Foraminifera and Radiolaria (Leckie et al., 2002). During the same period, we can find the rise of the Artostrobioidea (ca. 174 Ma), the divergence of the superfamilies Pterocorythoidea and Lychnocanoidea (ca. 134 Ma) or the continuous diversification of the lineage III. After the OAEs and during the Early Cenozoic (ca. 66 Ma to present) populations isolated from each other and probably thrived by the favourable conditions, start diversifying into the most recent families (Acanthodesmoidea, Theopiliidae, Cycladophoridae fam. nov., Lychnocanoidea, Pterocorythoidea and probably Northotripodiscinioidea). During the Cenozoic, other groups were also recovered from small populations linked to climate oscillations such as Foraminifera (Hallock et al., 1991) or calcareous Coccolithophores (Bown et al., 2004). Finally, during the late Cenozoic, the most recent family (Carpocaniidae) and groups (i.e. *Lipmanella* clade within Plagiacanthoidea) appear or diversify probably just before the opening of the Tasmanian gateway between Antarctica and Australia to form Antarctic Circumpolar Current.

### Environmental genetic diversity of Nassellaria

Little is known about the diversity and ecology of contemporary Nassellaria. Despite recent studies using plankton net data publicly available (Boltovskoy et al., 2010; Boltovskoy and Correa, 2016; Boltovskoy, 2017), most of the knowledge on this taxa is inferred from their extensive fossil record worldwide (Petrushevskaya, 1971a; De Wever et al., 2001). The present morpho-molecular framework proposes a morphological classification based on the overall structure, a feature easier to access compared to internal spicular structures, and therefore facilitating morphology-based ecological studies. Our framework also establishes the reference basis for sequences-based environmental diversity survey and molecular ecology studies, allowing the accurate and fast taxonomic placement of metabarcoding data.

Here we considered publicly available environmental sequences closely related to Nassellaria, from various environment (e.g. Edgcomb et al., 2011; Orsi et al., 2012; Lie et al., 2014) into our morpho-molecular data frame (Fig. 7). Most of environmental sequences clustered within the Superfamily Plagiacanthoidea, revealing an underrepresentation of this taxon in our study despite the number of sequences included (23%). This clade can be found at high relative abundances all year round (Motoyama et al., 2005; Ikenoue et al., 2015), in every latitude (Boltovskoy et al., 2010) and at every depth (Boltovskoy, 2017). Most of the known plagiacantoid species are very small related to the average nassellarian size, so they could escape plankton nets or be overlooked during isolation. Also, Acanthodesmoidea and Artostrobioidea are two clades highly represented in environmental sequences compared to the number of morphologically described sequences included in our study.

The former superfamily is represented by sequences coming mainly from the subtropical and tropical South China Sea where Radiolaria are very abundant (Wu et al., 2014). The Artostrobioidea is mainly characterised by deep environmental sequences, an environment poorly represented in our phylogenies mainly due to the difficulties in the DNA amplification of nassellarian specimens habiting the deep ocean. Likewise, the Superfamily Acropyramioidea, restricted to the deep ocean, is represented by only one sequence of the 28S rDNA gene. Therefore, due to the lack of 18S rDNA marker for this clade, it is likely that some of the sequences clustered within this lineage actually belong to the Acropyramioidea, since most of them come from deep environments (e.g. Kim et al., 2012; Lie et al., 2014; Xu et al., 2017). Increasing molecular coverage of the reference sequence database, through exploration of a variety of ecosystems, will be critical to properly decipher placement of environmental sequences. Our integrated approach enables providing a robust evolutionary history and classification of Nassellaria, and allows future ecological and diversity studies based in both a morphology or through a metabarcoding approach.

## Acknowledgments

This work was supported by the IMPEKAB ANR 15-CE02-0011 grant and the Brittany Region ARED C16 1520A01, the Japan Society for Promotion of Science KAKENHI Grand No. K16K0-74750 for N. Suzuki and “the Cooperative Research Project with the Japan Science and Technology Agency (JST) and Centre National de la Recherche Scientifique (CNRS, France) “Morpho-molecular Diversity Assessment of Ecologically, Evolutionary, and Geologically Relevant Marine Plankton (Radiolaria)”. We are grateful to T/S Oshoro-maru (Hokkaido Univ.), T/S Toyoshio-maru (Hiroshima Univ.) and their directors, Susumu Ohtsuka (Hiroshima Univ.) and Atsushi Yamaguchi (Hokkaido Univ.), as well as Akihiro Tuji, Yasuhide Nakamura and Yoshiaki Aita for sampling in Oshoro-Maru. We would like to thank the MOOSE cruise and program for the opportunity of sampling and the facilities given onboard, as well as John Dollan for hosting us multiple times in the Laboratoire d’Océanographie of Villefranche sur Mer. We thank Sebastien Colin for the Confocal Microscope imaging. We are also very grateful to Peter Baumgartner for his valuable comments in geological events, and specially to Luis O’dogherty, Spela Gorican and Taniel Daenelian for sharing their knowledge in the fossil record and improving the calibration of the molecular clock.

## Taxonomic Notes

Following the molecular phylogenetic results, new suborders are established here, and the definition of superfamilies is revised here following ICZN2000.

Suborder **Multicyrtidina** Pessagno, 1977 stat. nov. Reference superfamily: Eucyrtidioidea.

Definition: This suborder is relevant to lineage I. Multi-segmented nassellarians with a simple cephalis. Segmentation is divided by inner rings or relevant structure.

Remarks. Suborder Multicyrtidina includes only one Superfamily Eucyrtidioidea in living radiolarians, but presumably include a large number of families and genera of extinct radiolarians. The word “multicyrtid” appeared first in Pessagno, (1977, P. 53), as far as known.

Superfamily **Eucyrtidioidea** Ehrenberg, 1846 sensu emend here by Suzuki.

Type genus. *Eucyrtidium* Ehrenberg, 1846 (type species: *Lithocampe acuminata* Ehrenberg).

Emended definition. A spherical cephalis with a sharply constricted basal aperture and many segmentations which are generally divided by inner rings. No feet.

Remarks. This Superfamily corresponds to clade A. Although many suggestions about the definition have been made so long (see Matsuzaki et al., 2015), these two points are only different points from other nassellarians.

Suborder **Hephaistocyrtidina**suborder nov. by Sandín, Not and Suzuki. Reference superfamily: Acanthodesmoidea.

Definition: This suborder is relevant to lineage II.

Remarks. Hephasitocyrtidina subord. nov. includes Acropyramoidea, Carpocaniidae, Artostrobiidae and Acanthodesmidae. Although no morphological commonality has been found yet, pseudopodia tend to be present in many members. No protoplasmic features are known for Acropyramoidea. The name is derived from the Greek God “Hephaistos” of variable astonishing skills.

Suborder **Vestiacyrdina**suborder nov. by Sandín, Not and Suzuki. Reference superfamily: Plagiacanthoidea.

Definition: This suborder is relevant to lineage III.

Remarks: Vestiacyrdina subord. nov. includes Northotripodisciniodea, Theoperidae sensu strict, and Plagiacanthoidea. Although morphological commonality has not been confirmed, the members of Vestiacyrtidina tends to have a common feature of the combination with a very small volume of protoplasm, fragile skeleton (*Northotripodiscinus* is an exceptional case), absence of radiated pseudopodia (*Pseudocubus* is an exceptional case), and two or reduced one segment(s). The name is derived from the Roman “Vesta” whose stature or symbol was widely distributed in every house as a god of family.

Superfamily **Nothotripodiscinioidea** Deflandre, 1972 stat. nov. by Sandín, Not and Suzuki.

Type genus. *Nothotripodiscinus* Deflandre, 1972 (type species: *Nothotripodiscinus johannismonicae* Deflandre).

Definition. Practically single segment, although the upper and lower parts can be recognized by the position of *MB*. Initial spicular system is characterized by a very short *MB* (or missing) to form a three-pointed star rod system and a significant circular frame. Three feet or relevant structure is present.

Remarks. This corresponds with our Clade X. The genus *Nothotripodiscinus* was used to be synonymized with *Archipilium* since Petrushevskaya (1975), but the figured specimens of the type species of *Archipilium* apparently lack a three-pointed star rod system and a significant circular frame”. The intermediate form between *Northotripodiscinus* and *Enneaphormis* was described as “*Enneaphormis trippula*” by Renaudie and Lazarus (2015).

Family **Nothotripodiscindiae** Deflandre, 1972.

Type genus. *Nothotripodiscinus* Deflandre, 1972 (type species: *Nothotripodiscinus johannismonicae* Deflandre).

Remarks. Here we include genera *Archibursa* Haeckel, 1882 (type species: *Archibursa tripodiscus*

Haeckel), *Chitascenium* Sugiyama, 1994 (type species: *Chitascenium cranites* Sugiyama), and *Enneaphormis* Haeckel, 1882 (type species: *Sethophormis* (*Enneaphormis*) *rotula* Haeckel), because they have a three-pointed star rod system and a significant circular frame.

Superfamily **Plagiacanthoidea** Hertwig, 1879 emended here by Sandín and Suzuki.

Type genus. *Plagiacantha* Claparede and Lachmann, 1859 (type species: *Acanthometra arachnoides* Claparède).

Emended definition. Test can be divided into upper and lower parts by the level of *MB* or the neck of test. Test is small, and the initial spicular system contains variable types of arches. By these arches, antecephalic lobe, eucephalic lobe and postcephalic lobes are formed. This Superfamily also includes nassellarians made of bony frame only.

Remarks. This definition includes both Plagiacanthoidea and Cannobotryoidea. The name

“Plagiacanthoidea” has a priority to Cannobotryoidea.

Suborder **Jatagarasucyrtidina** suborder nov. by Sandín, Not and Suzuki. Reference superfamily. Pterocorythoidea.

Definition. This suborder is relevant to lineage IV.

Remarks. Jatagarasucyrtidina subord. nov. is comprised with the family Cycladophoridae fam. nov., and superfamilies Lychnocanoidea and Pterocorythoidea. The common morphological structure is three wing-like rods or rims from *D*-, *Lr*- and *Ll*-rods, two segmentation, and distinctive cephalis and thorax boundary. The name is derived from a mythological bird “Yatagarasu” in Japan which has three legs.

Family **Cycladophoridae** fam. nov. Suzuki

Type genus. *Cycladophora* Ehrenberg, 1846 (type species: *Cycladophora davisiana* Ehrenberg)

Definition. Initial spicule with *A*-, *V*-, *D*- and two *L*-rods, and an *MB*. No *PR* and No tubular cephalis horn. Test robust, helmet-conical, consisting of two segments with or without frill-like fringe.

Cephalis small, spherical, and pore-less or relict pores; distinctive from thorax; three wing-like rod or rims on upper thoracic wall.

Remarks. This new family is newly proposed to separate genus *Eucecryphalus*, the type genus of the family Theopiliidae, from *Cycladophora* which was used to belong to the Theopiliidae. More study needs to determine differences between Cycladophoridae fam. nov. and Theopiliidae. The former’s test is robust whereas the latter’s one is fragile. Superfamily is not determined to Cycladophoridae because nothing is known about the phylogenetic relationship with the Mesozoic family Neosciadiocapsidae. Cycladophoridae fam. nov. differs from the Superfamily Lychnocanoidea in that the latter has three distinctive feet which are connected through thoracic rims from *A*-, *V*-, and two *L*-rods. Cycladophoraidae is easily distinguished from the Superfamily Pterocorythoidea in having a pterocorythid cephalic structure with special lobes.

## Supplementary Data

**Figure S1.** Molecular phylogeny of environmental sequences (18S rDNA) associated to Nassellaria. The tree was obtained by using a phylogenetic Maximum likelihood method implemented using the GTR + γ + I model of sequence evolution. PhyML bootstrap values (1000 replicates, BS) are shown at the nodes. Black circles indicate BS of 100% and hollow circles indicates BS > 90%. Branches with a double barred symbol are fourfold reduced for clarity.

**Table S1.**List of specimens used to obtain nassellarian phylogeny.

**Table S2.**List of environmental sequences related to Nassellaria.

